# Serum cytokines during acute respiratory infection and relationship to age

**DOI:** 10.1101/2025.09.29.679372

**Authors:** Aanya Gupta, Holden T. Maecker

**Affiliations:** Human Immune Monitoring Center (HIMC), Institute for Immunity, Transplantation, and Infection (ITI), Stanford University, Stanford, CA 94305

## Abstract

Influenza and other seasonal respiratory viruses affect millions annually. While age is generally known to be correlated with risk and outcome, the mechanisms underlying these differences, especially in the infected, have been poorly defined. Previous studies have focused primarily on cell-subset shifts in older adults, leaving a gap in understanding of how cytokine responses vary across the full age spectrum. Cytokine data from Luminex assays were compiled from Stanford clinical studies and after batch control filtering, the dataset included 181 healthy individuals (ages 4–97) and 27 symptomatic individuals (mostly confirmed influenza, ages 14–77). Ordinary least squares regression was applied to assess cytokine differences due to infection status, age, and their interaction, with statistical significance defined as p < 0.05. The results illustrated that pro-inflammatory cytokines were found to be significantly elevated in infected individuals, with a trend of stronger effects in the young supported by comparison of intercepts between the regressions for the healthy and infected cohorts. The concept of inflammaging was also seen through several biomarkers with significant non-zero slopes in healthy cohorts that were not well-established in prior research. Our findings reveal both expected and novel cytokine behavior in influenza infection across a wide range of ages. By incorporating younger individuals and including age as a continuous variable, the study makes progress towards a deeper understanding of the changes in the immune system and its response to influenza across the lifespan.

## Introduction

Respiratory viruses, including influenza, RSV, and coronaviruses, continue to be a large burden on global healthcare systems. Influenza, commonly known as the flu, severely affects 3-5 million people annually, and causes 290,000 - 650,000 deaths every year [1]. Although the reach of influenza is widespread, there are certain factors that cause some groups to be more susceptible than others. Pre-existing health conditions and immunocompetence are established determinants, since both suppress the immune system and can also increase the severity of the infection [2].

Another significant component in an individual’s response to influenza is age. The elderly are at a higher risk of severe outcomes and mortality, especially with novel strains of influenza, while the young have higher infection rates and hospitalization [3]. Still, the causes for these differences between the immune systems of young and old are not clearly defined. Older immune systems suffer from immunosenescence, with a weakened humoral and cellular immune response [4]. Seniors are more likely to have reduced naive T and B cell production and impaired function of many key types of immune cells [4]. On the other hand, the young, with less immune memory, rely on innate immunity, meaning that there are increased naive T and B cells but also the possibility of hyper-reactive innate responses [5]. Another key difference is the impact of inflammaging, where immune dysregulation and elevated baseline cytokine levels drive chronic inflammation with age [6]. Microbiome composition has also been theorized to influence the imbalance of cytokines, as young microbiomes are still developing and older microbiomes become less diverse and more inflammatory [7].

Investigation of changes in cell proportions in the young versus old when infected has already been explored. Previous research has found that significant changes in cell subset frequencies are observed mostly in young individuals [8]. Larger shifts are seen only in the youth, although the elderly did show baseline proportions that aligned with expectations of aging’s effect on the immune system. Ugale et al. theorize that the older cohort’s more static immune reactions are a result of both immunosenescence and earlier exposure, which allows their systems to recognize the infection when exposed to influenza in their later years [8].

However, there is not much exploration of other immune system components in response to respiratory infections at different ages. Analysis of the trends in cytokine data of individuals’ response to influenza with respect to age has been limited. Filling this gap would allow us to answer the question of whether cytokine responses exhibit similar age-associated patterns. Thus, in this paper, we examine these cytokine data from the Luminex assay through statistical analysis of both healthy and influenza-infected subjects with diverse ages. Using the Mann-Whitney U-Test and regression analysis to separately identify significant differences in cytokine levels due to age and infection, we develop a more comprehensive understanding of how immune function evolves across the lifespan in the context of acute respiratory infection.

## Materials and Methods

### Study subjects

The study dataset was created with two cohorts, infected and healthy. The infected cohort was from study SLVP022, consisting of individuals presenting in an outpatient clinic with acute respiratory infection. 79% of these were confirmed to be infected with influenza A or B [8]; but all subjects were retained in the study, since those who were negative might have been undetected due to strain heterogeneity or low viral load. Age-matched control subjects were chosen from concurrent studies of influenza vaccination in healthy individuals (studies SLVP015, SLVP021, SLVP028, SLVP029, SLVP030, and SLVP031). All studies were approved by the Institutional Review Board at Stanford University.

### Data selection

To avoid confounding effects of different Luminex kit batches, only data from a single Luminex batch (H63, 01-27) were used, ending in a dataset of 27 people with ages ranging from 14 to 77 years old. The same filtering was done for the healthy cohort, resulting in 181 individuals with ages ranging from four to 97 years old. **Table 1** summarizes the number of subjects and range of ages for each study that were used for the present analyses.

**Table 1.**
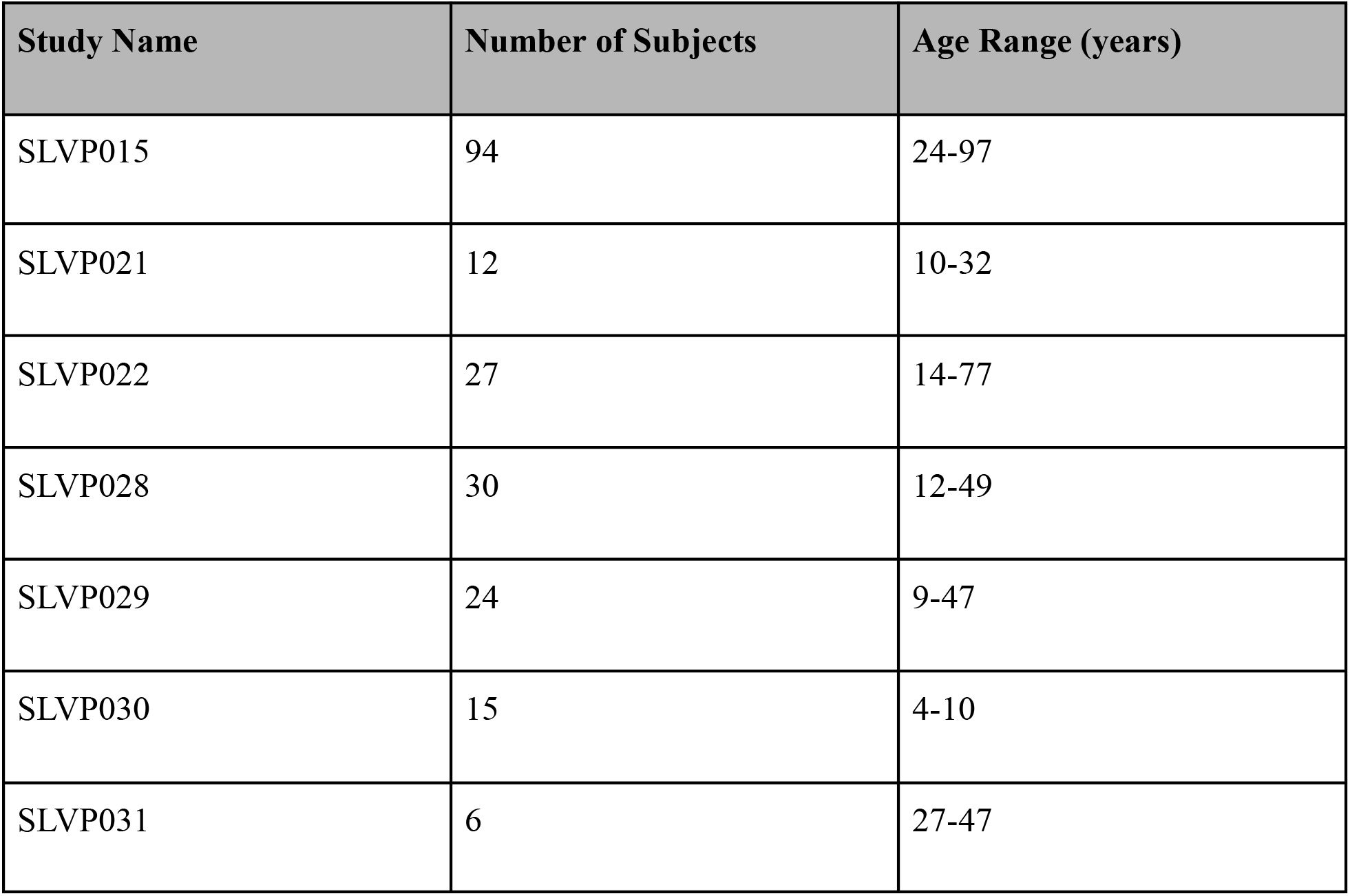
Description of Included Influenza Studies.

| Study Name | Number of Subjects | Age Range (years) |
| --- | --- | --- |
| SLVP015 | 94 | 24-97 |
| SLVP021 | 12 | 10-32 |
| SLVP022 | 27 | 14-77 |
| SLVP028 | 30 | 12-49 |
| SLVP029 | 24 | 9-47 |
| SLVP030 | 15 | 4-10 |
| SLVP031 | 6 | 27-47 |

### Controls and batch effects

To identify any possible remaining batch effects, the shared serum control samples from each included assay were compared. These consisted of one of two different controls, labeled CONS2 and CONS3. For both, the coefficient of variation (CV) of the combined set of their values from all assays in each study were calculated. The CV for CONS2 was found to be 16.5%, and for CONS3, 9.4%. These relatively small CVs provide evidence that persisting batch effects were minor, and suggesting that more significant variation among the subjects was due to age or infection status.

### Statistical analyses

Regression analysis was performed with both cohorts. For each biomarker, the subjects were plotted with age as a continuous variable on the x-axis and the value measured for the biomarker on the y-axis. Best-fit lines were found for each cohort using ordinary least squares (OLS), with the slope and intercept (the coefficients of the equations) for both lines being recorded. The p-values, which were calculated from a t-test with the standard errors derived from the residuals of the line, were used to determine whether the differences in the slope and intercept between the groups were statistically significant. For each biomarker, they were adjusted using the False Discovery Rate Benjamini-Hochberg procedure (threshold of adjusted p-value < 0.05). The p-values were also used to determine whether the slope of the best-fit line of the healthy cohort was significant, to ensure that the data aligned with the expected change in cytokine baseline levels simply due to age. Another linear regression model (OLS) was applied with biomarker values as the dependent variable and infection status as the independent variable, without adjusting for age. Similarly calculated p-values were used to determine whether the difference in distributions of the biomarker levels for both groups was significant (p-value < 0.05).

## Results

### Cytokine differences with infection

Fitting a linear regression model with the biomarker values and infection status allowed us to assess whether infection status alone was associated with differences in biomarker levels. **Figure 1** depicts the distributions of values for the four cytokines that have the most significant differences in these distributions of values between the two cohorts of the dataset. These include IP-10 (p=1.26E-11), RANTES/CCL5 (p=0.000283), IL1RA (p=0.00407), and IL7 (p=0.00451). (**Supplementary Table 1** contains the seven remaining biomarkers with significant differences and their p-values). The levels in the infected group were found to be higher for all the above markers except IL7.

**Figure 1.**
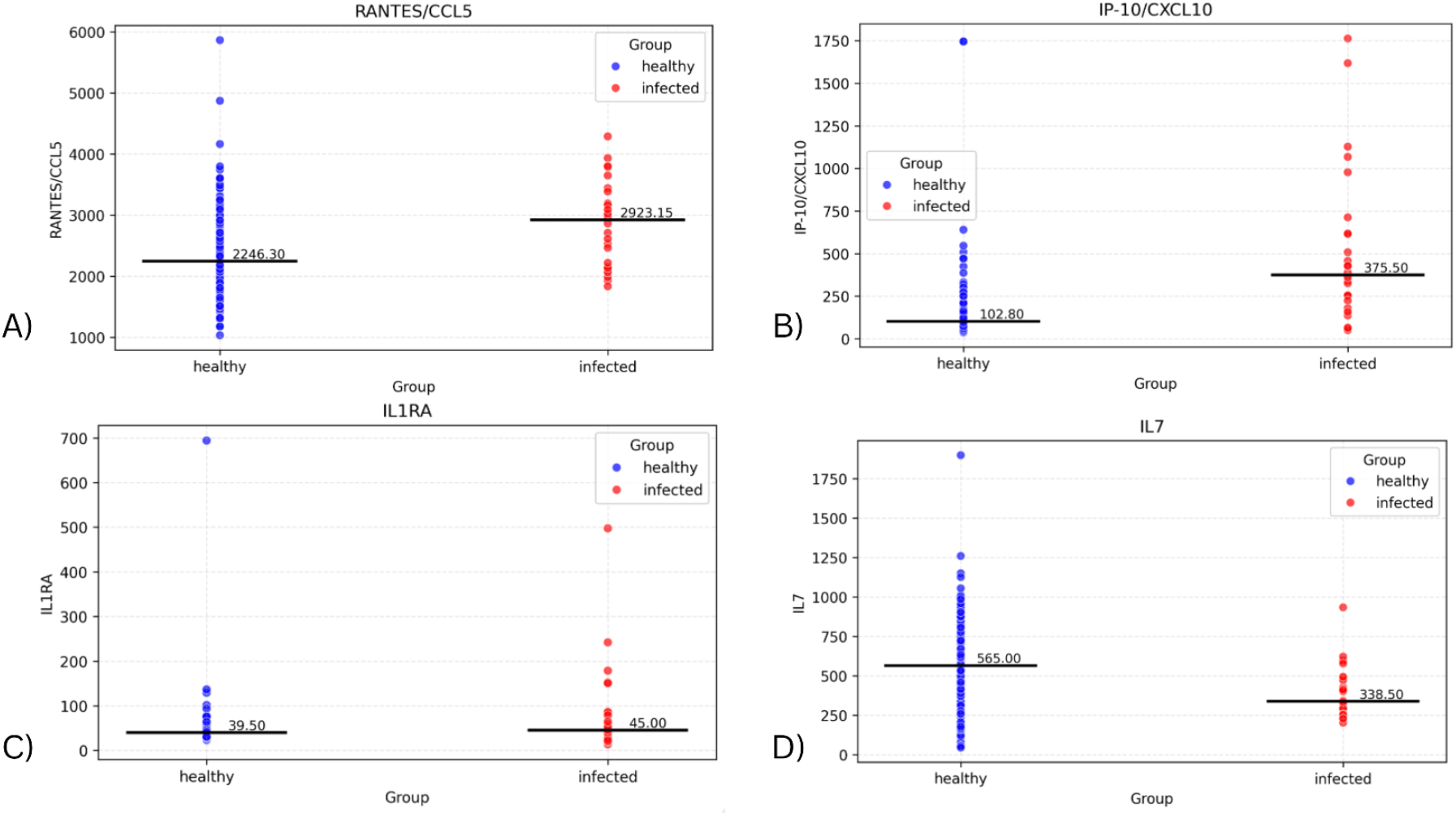
Plots showing distribution of subjects for biomarkers with significant difference in values between the infected and healthy cohorts.

### Effect of age on cytokine differences with infection

To assess whether the effect of infection on cytokine levels differed by age, a linear regression model was fitted on each cohort, with the interaction of age with biomarker and group also considered. Significant differences in the slope of both regression lines were found in four biomarkers: GROA (adjusted p=0.00712; p=0.00356), ENA-78 (adjustedd p=0.00460, p=0.00230), IL31 (adjusted p=0.00781; p=0.00391), and IL18 (adjusted p>0.05; p=0.0421) (**Figure 2**). Comparing the slopes of these regression lines identifies any effects of infection on the change in biomarker values with age. In all four cytokines, the infected slope was found to be higher than the healthy, signifying a more pronounced response to infection in the elderly. However, the infected slope appeared to be highly influenced by a single elderly outlier, suggesting that these differences may not represent a consistent biological trend.

**Figure 2.**
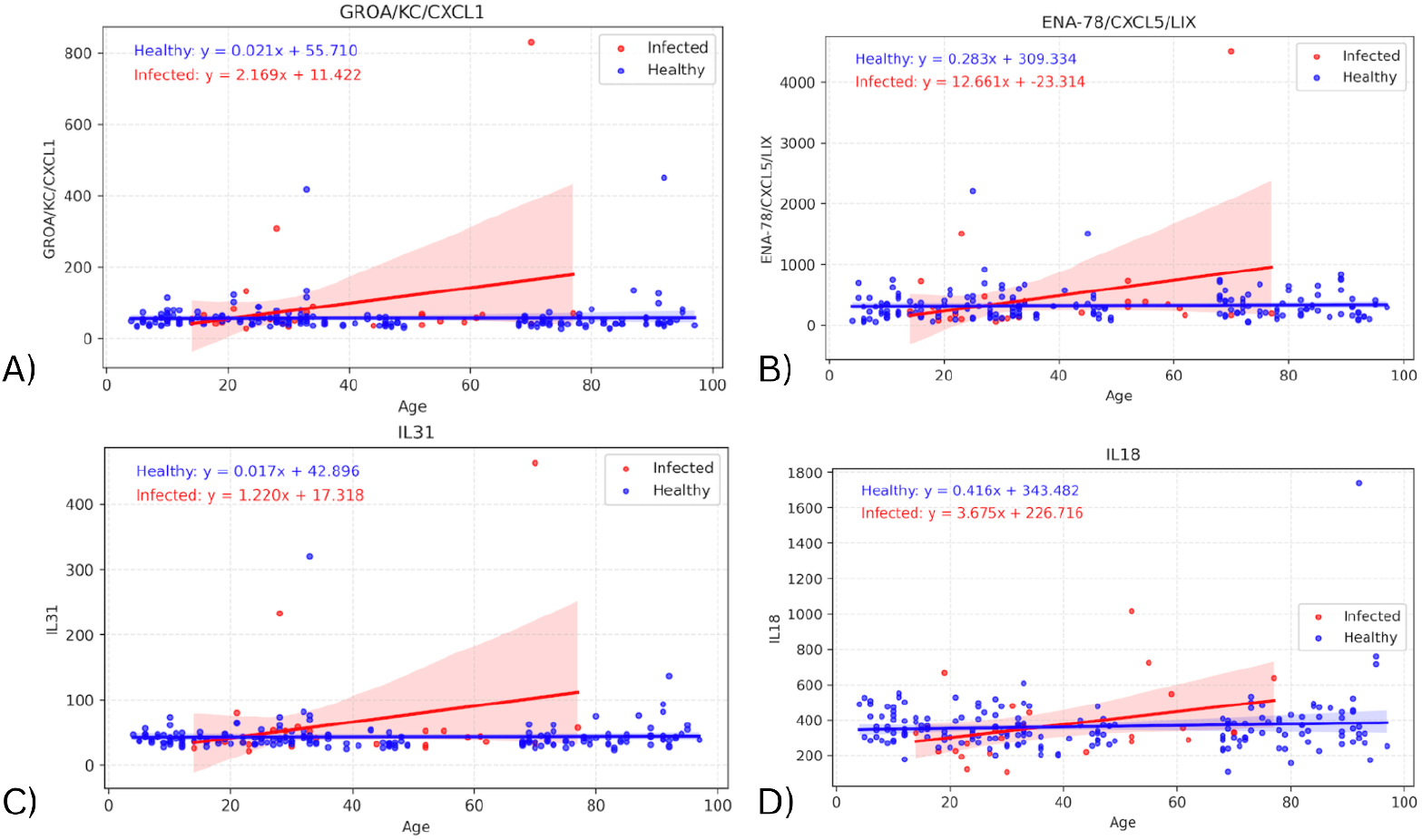
Graphs showing regression lines for all biomarkers that have significant difference in slopes between infected and healthy cohorts.

The same linear regression models were used to identify significant differences in intercept between the infected and healthy groups. These were only found in two biomarkers: IP-10 (p=0.00129) & RANTES (p=0.00653), shown in **Figure 3**. For these cytokines, the intercept difference suggests an effect of infection on the youngest individuals, as the intercept represents the predicted cytokine value at birth (age 0). In the graph for RANTES, the difference in biomarker values appears to decrease as age increases (though the slope difference was not significant). For IP-10, the slopes for infected and healthy groups appear very similar, suggesting a consistent infection effect regardless of age.

**Figure 3.**
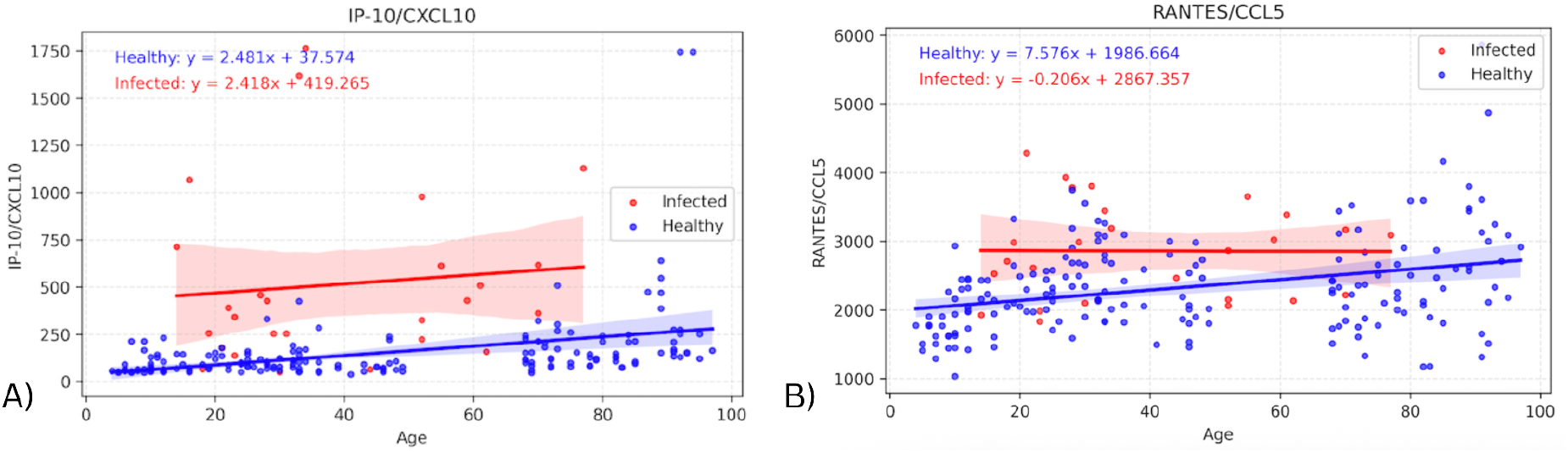
Graphs showing regression lines for all biomarkers that have significant difference in intercepts between infected and healthy cohorts.

### Cytokine differences due to age alone

The slope of the healthy cohorts for each cytokine were also investigated on their own, with the p-value from the linear regression being used to determine whether the slope was significantly non-zero. **Figure 4** depicts the regression lines for the four biomarkers with the most significant healthy slopes: EOTAXIN (p=2.29E-09), LEPTIN (p=1.78E-07), MCP1 (p=1.19E-07), and VCAM1 (p=1.45E-06) (**Supplemental Table 2** contains the remaining seventeen cytokines and their corresponding p-values). These biomarkers support the expected change in levels due to age, and also indicate an effect due to infection that is separate from the effect of age on cytokine values. In particular, the regression lines in Figure 4 show clear upward slope in the healthy cohort, consistently positive across all four biomarkers. The infected cohort has greater variability in slope (though it is still positive), suggesting that while age-related increases are apparent in healthy individuals, infection introduces additional dispersion in cytokine levels.

**Figure 4.**
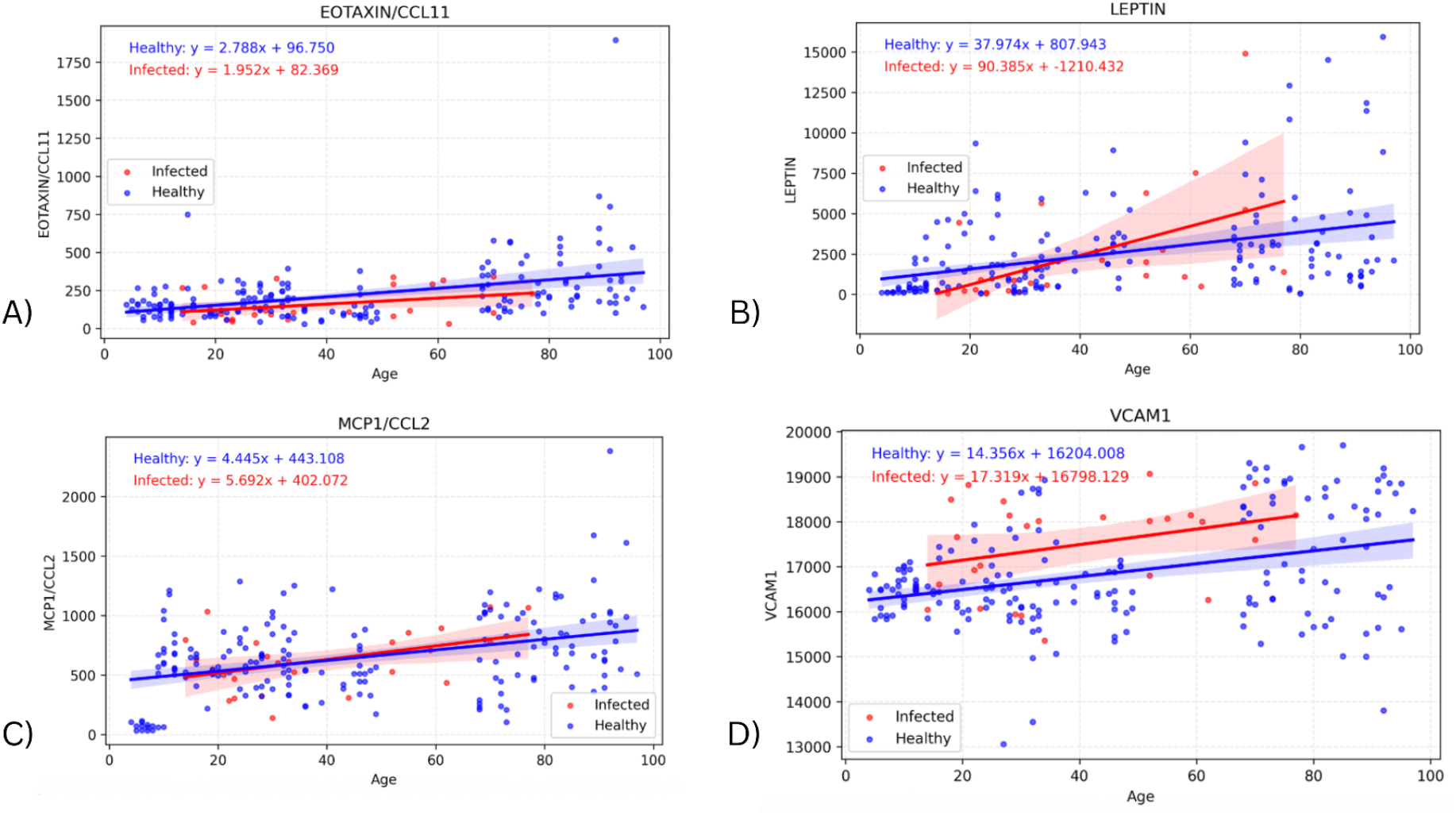
Graphs showing regression lines for the top four biomarkers that have a significant non-zero slope for the regression of the healthy cohort.

## Discussion

Through our analysis, we explored the relationship between respiratory infection and serum cytokine levels in individuals of diverse ages. A total of 11 of 63 measured cytokines were differentially expressed in infected vs. healthy individuals. Except for IL-7, all of these were elevated in the infected group. While the elevated cytokines were primarily pro-inflammatory, there were notable absences, such as IL-6 and IFNg, which were not significantly different. Such cytokine elevations have previously been associated with the presence and severity of influenza-associated illness [9, 10]. We may have observed fewer and smaller cytokine elevations in our study, as our subjects were in an outpatient setting, rather than hospitalized. On the other hand, our more comprehensive panel of 63 cytokines may have revealed elevations in cytokines not tested in these previous studies.

The significant Y-intercept differences for IP-10 and RANTES suggest that these chemokines are upregulated with infection in the youngest individuals (Y-intercept = age 0). While the elevation for IP-10 in infected individuals appeared to be similar across ages, that for RANTES appeared to decrease with age (though the infected slope was not significantly different from the healthy slope). Those cytokines with significantly different slopes with age in infected vs. healthy individuals all had higher infected slopes, i.e., greater elevations for older infected individuals. However, these seemed to be driven by one or two high outliers in the elderly population, so may not represent consistent findings.

These findings are in some contrast to those of our lab’s previous study of cell subset changes with influenza infection [8]. In that previous study, significant cellular changes with infection were confined to the young cohort, with no significant changes in the elderly. This reflected a trend towards more effector-like cell phenotypes in the elderly, which more closely resembled the young infected phenotype. In the current study, cytokine changes appear to be largely independent of age, or even greater in the elderly (with the caveat about outliers mentioned above). One possible exception was RANTES, for which elderly infected individuals seemed to be more similar to healthy; but the slope differences with age between healthy and infected subjects were not statistically significant. All in all, this suggests that, while the cellular immune system appears to adapt with age and show smaller changes with infection in the elderly, cytokine responses tend to be relatively consistent across ages.

We also found 21 cytokines showing significant positive slopes with age in healthy individuals. These increases with age may be related to “inflammaging”, as older immune systems experience more cytokine dysregulation [11]. However, the cytokines identified as age-associated in our study have generally not been so associated in previous studies.

An earlier investigation found levels of TNFɑ, IL-17, IL-6, IL-12p70, IL-13, IL-10, and IL-17 to be highly correlated with age [12]. These discrepancies may be due to our inclusion of a broad age range, including younger samples. Prior work has largely found changes in the most elderly healthy adult subjects, e.g., seeing significant shifts in cytokine levels only above 86 years [13].

Together, our findings present both expected and novel aspects of cytokine regulation in the context of aging and influenza infection. While increases in pro-inflammatory cytokines align with their known antiviral roles, we also found cytokine differences with infection not previously emphasized in influenza studies, perhaps in part due to the large panel of cytokines we investigated. Most cytokines that were differentially expressed with infection were not reliably age-associated, in contrast to our previous findings with cell subset changes with infection, which were all decreased in the elderly. Together, these results highlight the importance of studying immune responses across the full age spectrum.

Our study’s major limitation was the relatively small size of the infected cohort (n=27). With more individuals, especially at the extremes of age, we could become more confident of the significant findings with regard to slopes with age. Another caveat is potential batch effects, which were confounded by the fact that healthy and infected subjects were run in separate experiments. We feel this issue was minimized by the use of a single batch of Luminex kits across all experiments, and the fact that shared controls had a relatively low C.V. across experiments. As such, we hope this study will inspire more detailed analyses of cytokines and their role in acute respiratory infections across the spectrum of age.

## Supplemental Figures

**Supplementary Table 1** - List of all biomarkers with significant difference among distribution for healthy and infected cohorts

**Supplemental Table 2** - Cytokines with significant slopes among the healthy subjects

## Acknowledgements

The authors gratefully acknowledge Cornelia Dekker, Mark Davis, and Harry Greenberg for clinical study design and execution for all cohorts. We thank Yael Rosenberg-Hasson and Iris Herschmann for performing the Luminex assays, and Janet Siebert and Wes Muncil for creating the Stanford Data Miner that allowed for integration of the data.

